# Autoencoders for unsupervised analysis of rat myeloarchitecture

**DOI:** 10.64898/2026.03.02.708929

**Authors:** Melina Estela, Raimo A. Salo, Isabel San Martín Molina, Omar Narvaez, Ville Kolehmainen, Jussi Tohka, Alejandra Sierra

**Affiliations:** A.I. Virtanen Institute for Molecular Sciences, University of Eastern Finland, Kuopio, Finland; Department of Applied Physics, University of Eastern Finland, Kuopio, Finland

**Keywords:** Histological image analysis, convolutional autoencoders, tissue clustering, image clustering, myelin, traumatic brain injury, neuropathology, computational pathology, unsupervised deep learning, Gaussian mixture models

## Abstract

Quantitative assessment of brain histology is often constrained by predefined feature sets and labor-intensive manual annotations. To overcome these limitations, we employed unsupervised deep learning to automatically extract and quantify tissue organizational patterns from myelin-stained rat brain sections without the need for prior labeling. We evaluated nonlinear convolutional autoencoders (AEs) against linear principal component analysis (PCA) for feature representation, followed by clustering with Gaussian mixture models. Compared to PCA, AEs better preserved fine axonal architecture and produced more consistent and interpretable tissue clusters across hierarchical levels. The resulting clusters revealed anatomically meaningful tissue organization, including different white matter densities and grey matter subregions. When applied to tissue from sham and mild traumatic brain injury animals, AE-derived features also captured pathology-related alterations, such as white matter loss and injury-specific microstructural changes. These findings demonstrate that unsupervised deep learning can automatically characterize tissue organization at multiple scales and detect pathological changes, offering a scalable, unbiased approach to quantitative neuropathology.

## Introduction

Histological analysis of healthy and diseased brain tissue has traditionally relied on expert visual inspection of stained sections to assess tissue architecture and identify pathological patterns. Over the last decades, automated whole-slide scanners have begun transforming histology ^1–4^ by enabling high-resolution digitization of large numbers of tissue sections. While this development has increased data volume and quality, it has also increased the workload of neuropathologists and neuroscientists. Consequently, the growth in digital histology data has created both an opportunity and a necessity for computational methods^4,5^, marking a transition from purely qualitative visual assessment toward quantitative, algorithm-driven analysis.

A major limitation of current quantitative histology workflows is their dependence on visual morphological assessment^6,7^ and manual delineation of regions of interest. Although effective for targeted analyses, these approaches are time-consuming, subjective, and difficult to scale to whole-section or whole-brain datasets. To address these challenges, several open-source software, such as QuPath^8^, Fiji^9^, and others^10,11^, automate parts of the image analysis pipeline, including cell detection, staining-intensity quantification, and morphometric measurements across large tissue areas. However, these tools remain largely object-centric, relying on predefined features, parameter tuning, or manual input, yet they do not provide whole characterization of brain tissue sections^2^.

In this context, computational histopathology has advanced substantially through supervised deep learning models trained on large annotated datasets^1,12^. These models enable automated prognosis prediction^12,13^, segmentation^13,14^, and tumor classification^15–21^. Similar approaches can be applied to brain tissue, supporting cell detection^22^, segmentation of cellular and subcellular structures, cytoarchitectonic mapping^14^, and myelin analysis^23^. However, these supervised approaches remain constrained by their need for annotated training data and predetermined tissue classes, limiting their ability to capture subtle variations in brain tissue architecture^11^.

Unsupervised learning offers an alternative for discovering patterns directly from data without manual annotations. While unsupervised deep learning has been applied in non-brain tissues^24–26^, it has not yet been applied systematically to myelin-stained brain sections, where axonal orientations, fiber crossings, and cellular distributions form complex microstructural patterns. A central question in applying unsupervised methods to histological data is whether nonlinear featureextraction techniques provide meaningful advantages over classical linear approaches. Nonlinear methods, such as autoencoders^27^ (AEs), can capture complex biological structure but require more computational resources, while linear methods, such as principal component analysis (PCA) offer interpretability and efficiency.

In this work, we applied convolutional AEs to extract features from myelin-stained rat brain sections and used unsupervised clustering to identify distinct patterns of myeloarchitecture. We compared these nonlinear representations with those derived from PCA, evaluating both their ability to capture biologically meaningful structural detail and their computational efficiency.

## Results

### Unsupervised clustering workflow results in systematic myelin tissue classification

We developed a computational workflow (Figure 1) to quantify how patterns of myelinated axons are distributed across brain tissue sections. To capture these spatial patterns, we systematically extracted non-overlapping square tissue patches at two scales —128×128 pixels and 256×256 pixels— from intensity-normalized myelin-stained rat brain sections.

**Figure 1:**
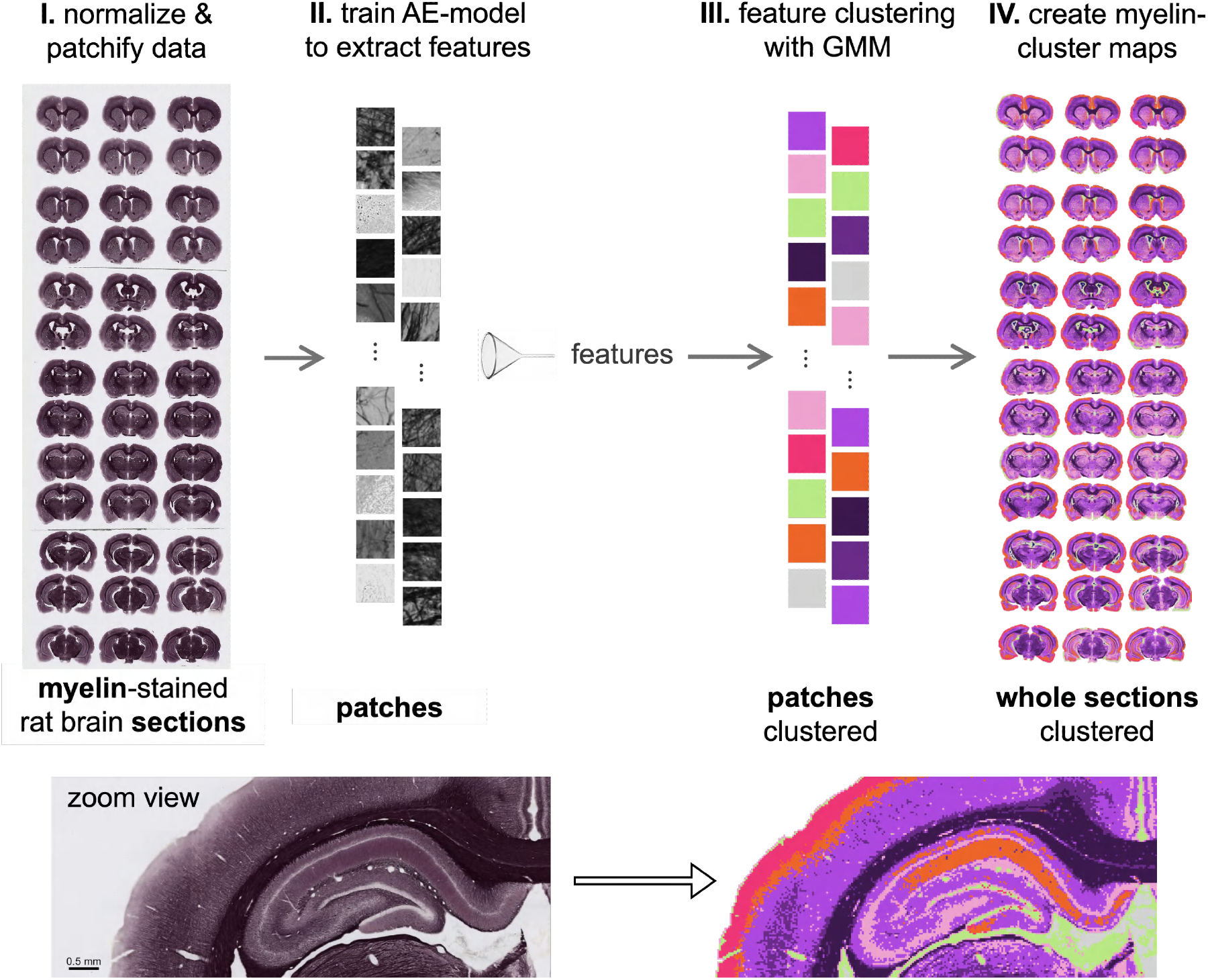
Workflow for feature extraction and classification of myelin histological rat brain sections. First, the myelin-stained rat brain sections are intensity-normalized and divided into non-overlapping patches at two scales: 128×128 pixels (model128) or 256×256 pixels (model256). A mid-coronal brain section is *∼*40,000×60,000 pixels, producing over 146,000 non-overlapping 128×128 pixel patches or over 36,000 256×256 pixel patches. Second, patches are used to train AE-models for feature extraction (256 features per patch). Third, the extracted features are clustered using Gaussian mixture models (GMM). Last, cluster assignments are projected back onto the tissue sections to create myelin-cluster maps. At the bottom, a zoom view of original tissue and corresponding cluster assignments.

We then trained convolutional AEs to compress each patch into 256 latent features and compared their performance against PCA for dimensionality reduction. All models were trained on patches from 9 rat brains (4 sham-operated and 5 with mild traumatic brain injury (mTBI); in total 420 sections total) and validated on 4 independent rat brains (2 sham-operated and 2 with mTBI; in total 194 sections). Results presented below are from the validation set.

To automatically separate tissue organization patterns, we clustered the extracted features using Gaussian mixture models (GMM) ^28^, identifying distinct tissue clusters based on myelin properties.

### AEs preserve structural detail better than PCA despite higher reconstruction error

We reconstructed tissue patches from the extracted features to assess feature quality for each method (Figure 2). AE model reconstructions preserved thin axonal structures and tissue texture more accurately than PCA. PCA reconstructions smoothed fine details but maintained overall tissue organization. When comparing input patch size, model128 reconstructed fine axonal features better in both PCA and AE. In contrast, model256 captured broader structural patterns but lost small-scale detail due to its higher compression ratio. AE reconstructions were more faithful in preserving myelinated axon patterns.

**Figure 2:**
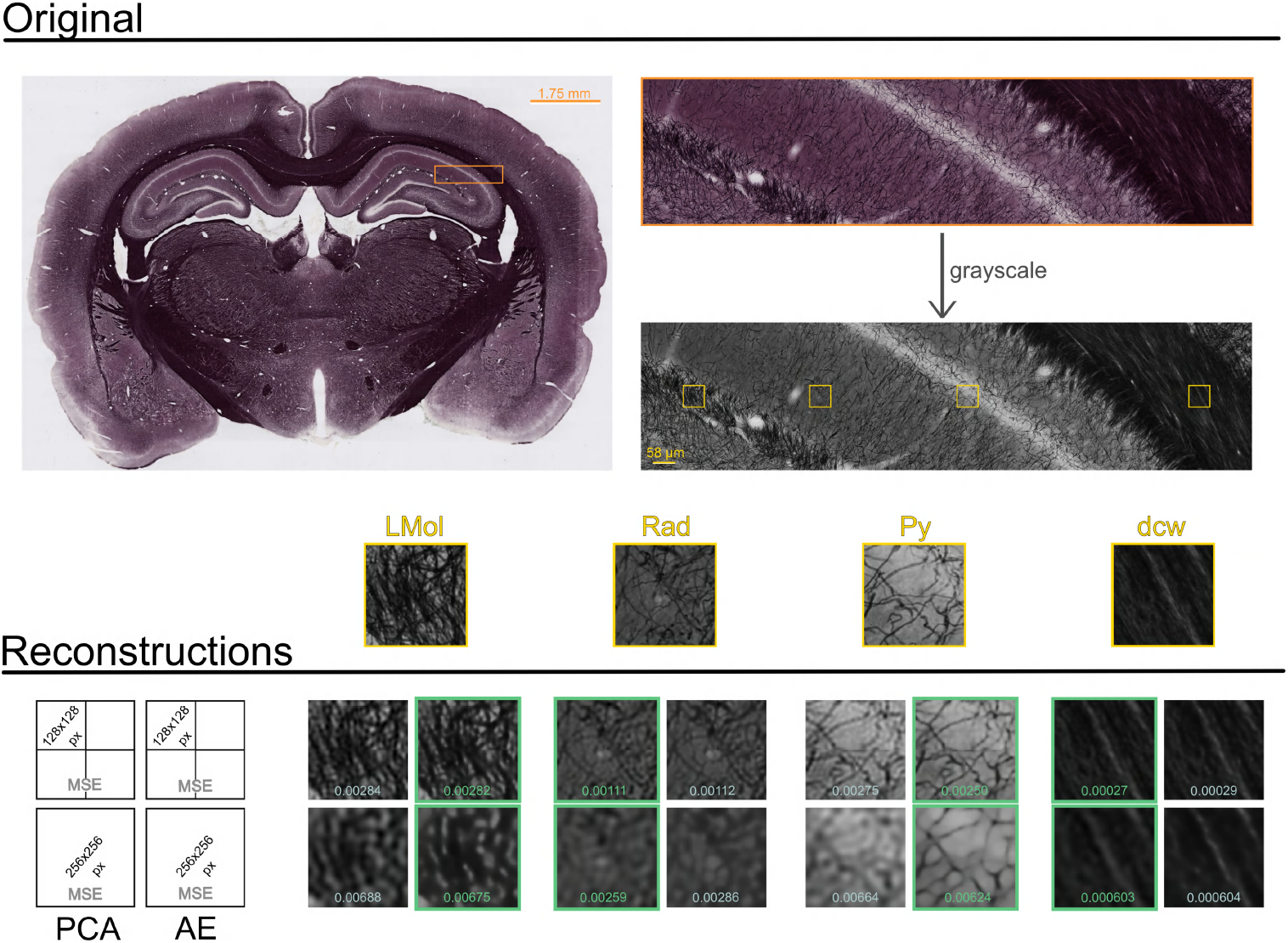
Original and reconstructed myelin-stained tissue patches. Representative mid-coronal section of a myelin-stained section of a sham-operated rat. Four tissue windows (58×58*µm*, 256×256 pixels) were reconstructed using four models: PCA-based and AE-based of both model128 and model256. Mean square error (MSE) values are included for each window, with green borders highlighting the better-performing model in each pair. Anatomical regions are: LMol (lacunosum molecular layer of the hippocampus), Rad (radiatum layer of the hippocampus), Py (pyramidal cell layer of the hippocampus) and dcw (deep cerebral white matter)^29^.

We assessed the reconstructions quantitatively through mean square error (MSE) and structural similarity index measure^30^ (SSIM). Individual patches showed highly similar MSE values between PCA and AEs, with each method outperforming the other on different tissue regions (Figure 2).

For MSE averaged across animals’ sections, PCA achieved slightly lower values than AE for both patch sizes (Table 1 and Table 2). For model128, the difference was negligible (Cohen’s d = -0.065). For model256, the effect remained small (Cohen’s d = -0.197).

**Table 1:** Reconstruction quality comparison for model128. MSE and SSIM values for PCA and AE across four validation animals. MSE values are scaled by 10^−3^ for readability. Lower MSE and higher SSIM indicate better reconstruction. Cohen’s d shows negligible effect sizes, with PCA slightly lower in MSE and AE slightly higher in SSIM. Bold values indicate the better-performing method for each metric.

| model128 | MSE ($\times 10^{-3}$) | | SSIM | |
| --- | --- | --- | --- | --- |
|  | PCA | AE | PCA | AE |
| sham 1 | 2.160 | 2.243 | 0.6914 | 0.6909 |
| sham 2 | 1.923 | 1.954 | 0.6863 | 0.6894 |
| mTBI 1 | 1.972 | 2.042 | 0.6930 | 0.6918 |
| mTBI 2 | 3.414 | 3.411 | 0.6568 | 0.6666 |
| mean | **2.367** | 2.412 | 0.6819 | **0.6847** |
| std | 0.705 | 0.677 | 0.0170 | 0.0121 |
| Cohen’s d | −0.065 |  | −0.189 |  |

**Table 2:**
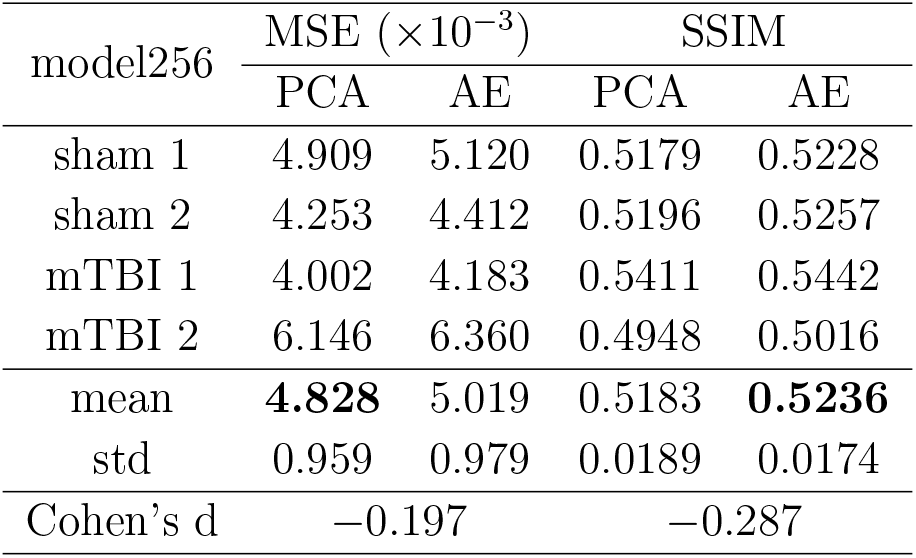
Reconstruction quality comparison for model256. MSE and SSIM values for PCA and AE across four validation animals. MSE values are scaled by 10^−3^ for readability. Lower MSE and higher SSIM indicate better reconstruction. Cohen’s d shows small effect sizes favoring PCA for MSE and AE for SSIM. Bold values indicate the better-performing method for each metric.

For SSIM, the pattern reversed, favoring AE with modest effect sizes. For model128, AE achieved higher SSIM than PCA, though the effect size was small (Cohen’s d = -0.189). For model256, the advantage for AE was more pronounced (Cohen’s d = -0.287). These findings suggest that while PCA achieved slightly better pixel-wise reconstruction accuracy, AE better preserved structural features, particularly when processing larger patches that capture more complex spatial patterns.

PCA models were trained in less than 3 CPU days, while AEs training required 6 GPU days for model128 and 9 GPU days for model256. AEs training used early stopping, halting when the validation error stopped improving for 10 consecutive iterations. Model128 achieved lower training and validation errors than model256 (Supplementary Figure 8).

### Both PCA and AE features enable identification of anatomically distinct tissue clusters

We performed unsupervised clustering using Gaussian mixture models (GMM) on the extracted features from both PCA and AE models to evaluate whether each model feature classified the tissue into biologically meaningful groups. We selected three, nine, and 21 as the number of clusters based on Bayesian Information Criterion (BIC)^31^ analysis (see Supplementary Figure 9 and Supplementary Figure 10), which showed an elbow at nine clusters with continued improvement with increasing number of clusters, suggesting different levels of tissue organization across multiple scales.

Figure 3 shows the clustering results from model128. When the number of clusters was three, both PCA and AE revealed similar broad anatomical divisions: highly myelinated areas (white matter, thalamic, and hippocampal areas); grey matter areas (thalamic, hippocampal, and cortical areas); and low intensity areas (borders with low staining intensity, ventricles, and the granule cell layer of the dentate gyrus). When the number of clusters was nine (Figure 3 second row), both methods resolved in more refined tissue structures including white matter, hippocampal subfields, and cortical layers. Differences in spatial organization, particularly in the cortical layers, began to emerge between the two feature extraction approaches. When the number of clusters was 21 (Figure 3 third row), both PCA and AE features enabled detailed tissue characterization. White matter was separated into clusters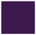,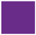 and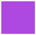 by both methods, reflecting variation in myelin density. In the cortex, AE-based features achieved clearer layer separation with clusters 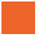,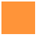 and 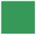, whereas PCA-based clustering showed less clear stratification in this region with clusters 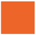, 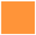 and 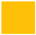.

**Figure 3:**
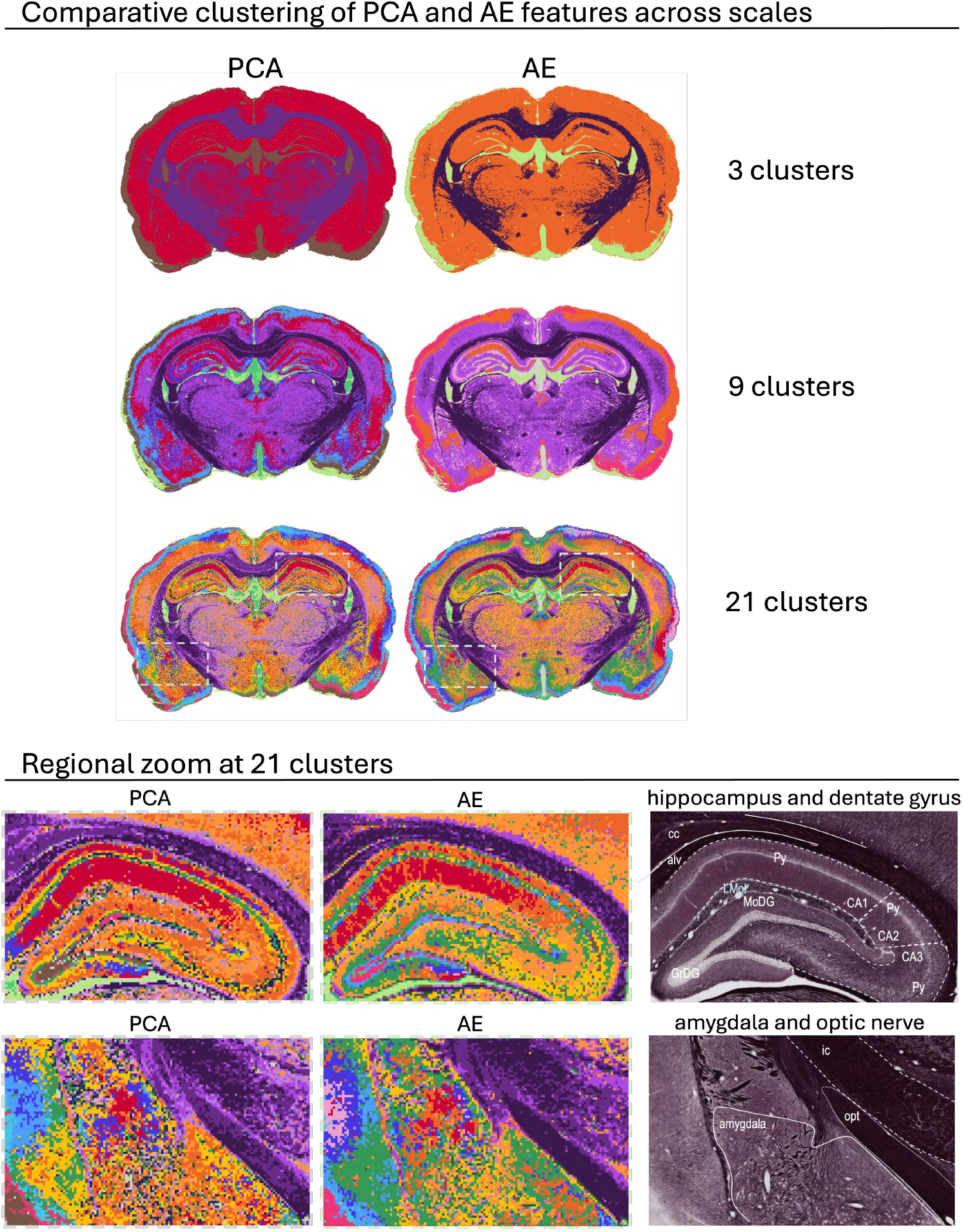
PCA versus AE feature clustering. Upper panel shows cluster assignments at three, nine and 21 clusters for PCA (left) and AE (right) features of the same section. Lower panel shows magnified hippocampal and amygdala regions at 21 clusters: PCA clustering, AE clustering, and the original myelin staining. Cluster colors were assigned at 21 clusters and matched for lower cluster numbers via symmetric KL divergence. Anatomical abbreviations in alphabetical order: alv, alveus of the hippocampus; CA1, field CA1 of the hippocampus; CA2, field CA2 of the hippocampus; CA3, field CA3 of the hippocampus; cc, corpus callosum; GrDG, granule cell layer of the dentate gyrus; ic, internal capsule; LMol, lacunosum moleculare layer of the hippocampus; MoDG, molecular layer of the dentate gyrus; opt, optic tract; Py, pyramidal cell layer of the hippocampus ^29^.

To visualize relationships across the three, nine and 21 cluster groups, we manually assigned colors to the 21 clusters. We then propagated these colors to the 9-cluster and 3-cluster solutions based on feature-space proximity, measured with the symmetric KL divergence. Interestingly, across different number of cluster groups, AE consistently assigned white matter to the same cluster 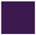, while PCA assigned white matter to different clusters depending on the total number of clusters. This suggested that AE captured white matter as a more compact, intrinsically distinct property.

A closer examination of the hippocampus and dentate gyrus at 21 clusters (lower panel of Figure 3) showed that the AE-feature-map had a clear cluster gradient of regions CA1, CA2 and CA3 of the hippocampus (with clusters 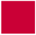, 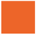, 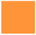), whereas the PCA-feature map showed a less gradual change. Both methods distinguished the lacunosum molecular layer, molecular layer of the dentate gyrus, and the granule cell layer of the dentate. The structure that PCA delimitated better was the pyramidal cell layer in the CA3. The zoom view in the amygdala and optic nerve showed also differences in the feature clustering. The optic nerve and internal capsule areas were clearly delimitated by clusters 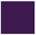,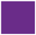 and 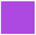 in the AE-map, while the PCA-map did not separate that neatly those regions. The AE-map also showed more compact delimitations of the subregions in the amygdala than the PCA-map.

Overall, AE-feature clustering showed greater stability across clusterings with different number of clusters and sharper anatomical boundaries than PCA-feature clustering. Subsequent results will therefore focus on AE-features.

### Centroids and probability maps enable anatomical interpretation of tissue clusters

We sought to understand what tissue properties each cluster represents and its spatial distribution. Figure 4 shows the probability maps of the 21-cluster map from the AE-features of model128, and the tissue patches whose latent space representation is closest to the cluster centroid. We found that in the mid-coronal section depicted in the figure, the white matter comprised 23.1% of total tissue content across three clusters (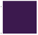, 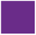, 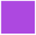): Cluster 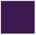 captured densely packed myelinated axons; Cluster 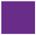 captured highly myelinated, oriented axons with inter-axonal spacing; Cluster 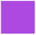 represented highly myelinated borders of major tracts (corpus callosum, internal and external capsules, optic nerves) with greater axonal separation.

**Figure 4:**
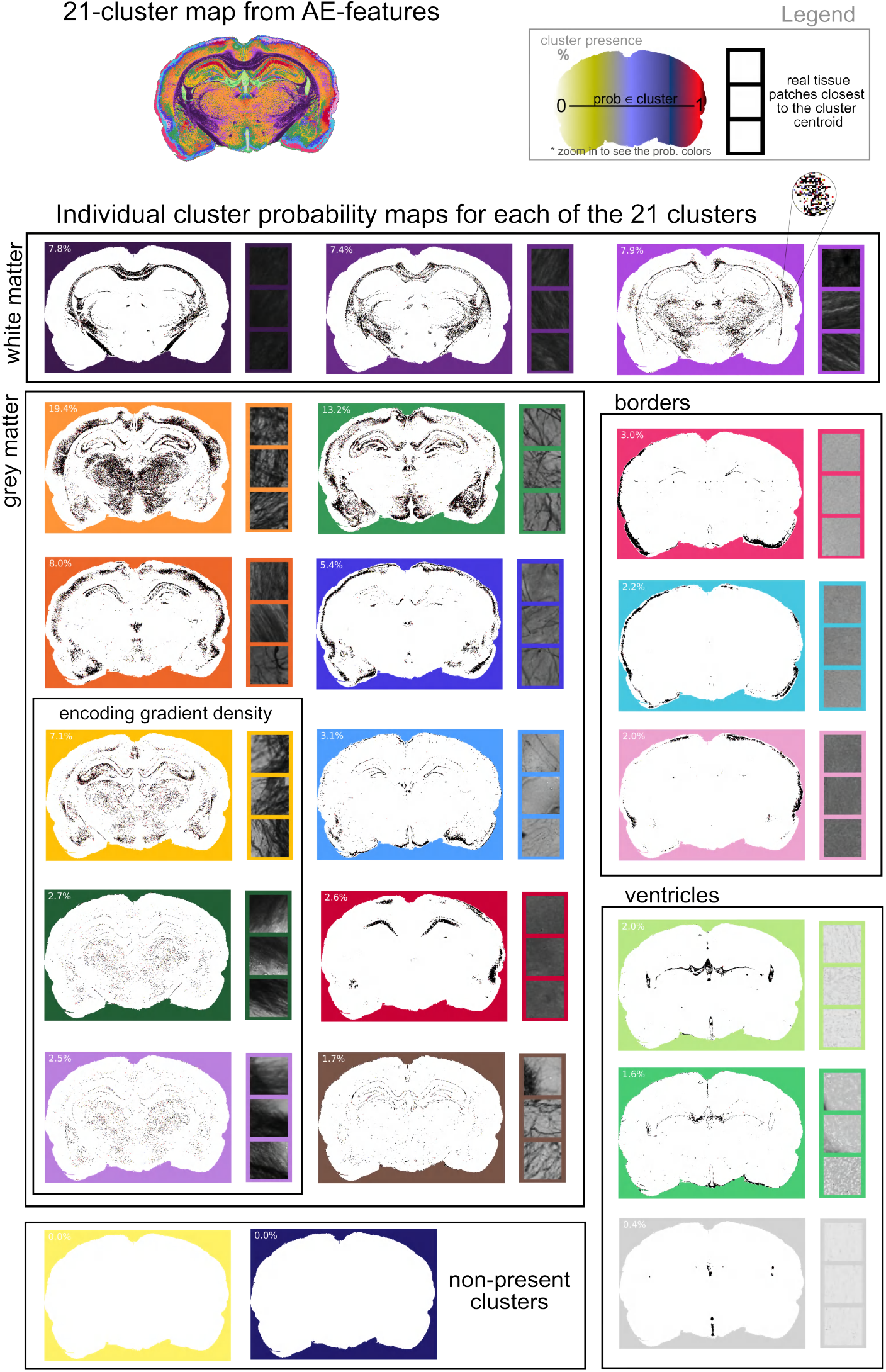
Probability maps of a single section. On the top left, overview of the 21-cluster map from AE-features from one section. On top right, the legend describing the color scale of the probability maps. Then, the individual cluster probability maps grouped in: white matter, grey matter, borders, ventricles and non-present clusters. Each probability map shows a percentage that indicates its presence in the tissue section, as well as three real tissue patches (128×128 pixels) closest to that cluster centroid.

Grey matter fell into 10 clusters. Cluster 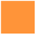 (19.4%) captured regions with abundant myelinated axons with varied orientations (with both aligned and crossing patterns) and visible interaxonal spacing. Cluster 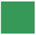 (13.2%) showed sparse, randomly oriented myelinated axons. Cluster 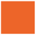 (8%) captured myelinated regions with loosely packed axons. Interestingly, clusters 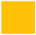,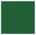,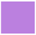 captured myelin-density gradients at boundaries between high and low myelin content.

Clusters 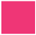, 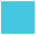 and 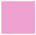 captured the section borders without myelinated axons but with varying tissue staining intensities. Three additional clusters captured ventricles and some tissue borders with staining artifacts. In this particular section, two of the 21 clusters were absent.

We observed no systematic differences between the left and right sides of the tissue section during clustering. This means that the resulting clusters were rotation-invariant although this was not explicitly enforced by the modeling scheme.

### Clustering identifies pathology-associated changes in white matter and myelin organization

To assess whether unsupervised clustering could capture tissue damage induced by mTBI, we analyzed myelin-stained sections from two sham-operated controls and two mTBI-injured animals (mTBI1 with subtle tissue damage and mTBI2 with evident tissue damage). We performed GMM on features extracted from patches across all four animals, enabling identification of common tissue properties and pathology-specific variations.

We present five representative clusters from the 9-component GMM model (probability maps shown in Figure 5). The first cluster delineated intact white matter, showing consistent presence in sham controls (*∼*13%) and mTBI1 (12%) but notably reduced in mTBI2 (9.7%), with striking asymmetry on the left (injured) side. Clusters A and B represented myelinated axons within grey matter regions at varying densities. In shams and mTBI1, clusters A and B represented >16% and >36% of tissue respectively; in contrast, these decreased to 8.8% and 27.3% in mTBI2. Most notably, cluster D captured border regions and hippocampal areas, with low presence in sham controls (*∼*5-6%) and modest elevation in mTBI1 (8.6%) but dramatic increase in mTBI2 (38.8%). This shift strongly suggested that cluster D specifically captured pathological tissue associated with mTBI.

**Figure 5:**
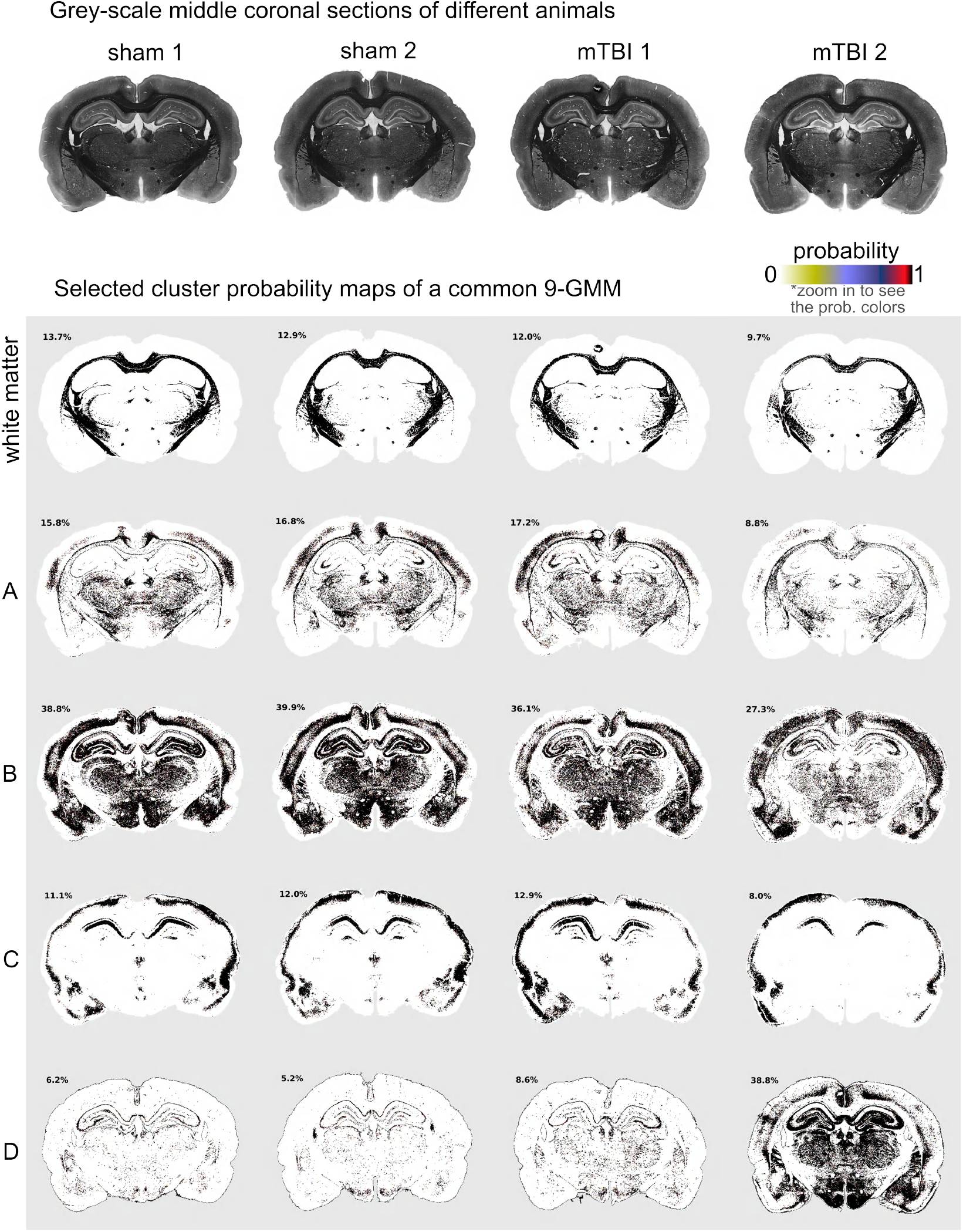
Representative cluster probability maps of a common 9-component GMM of four animals. Four representative coronal sections from four different animals (two sham and two after mild traumatic brain injury (mTBI)) and selected cluster maps from a common 9-component GMM.

## Discussion

Brain histology requires scalable methods that move beyond labor-intensive manual annotations and predefined features sets^12,32^. We proposed an unsupervised workflow that automatically quantifies spatial patterns formed by myelinated axons, enabling distinction of tissue organization and detection of pathological changes without requiring manual labeling of tissue classes or pathological features.

In the context of previous literature, our findings extend the use of unsupervised deep learning in histology. While prior studies have applied similar methods to non-brain tissues, such as H&Estained colon^24^, pancreatic^25^, or breast and lung tissues^26^, myelin-stained brain sections present unique challenges. White matter contains densely packed, overlapping myelinated axons with varying orientations, while grey matter exhibits lower but highly variable axonal density with diverse crossing patterns In regions like the cortex, these patterns transition gradually without clear boundaries. Successfully characterizing such tissue requires methods that preserve fine axonal details while capturing broader organizational features.

Our workflow extracts features from patches of myelin-stained rat brain sections and clusters these features to identify distinct tissue patterns. To evaluate feature extraction performance, we compared two dimensionality reduction methods: convolutional AEs and PCA. Both methods successfully extracted features capturing anatomical properties. However, AE outperformed PCA in preserving fine structural details of myelinated axonal tissue, and that MSE did not necessarily reflect preservation of visually important biological features.

When clustering the extracted features using GMM, we recovered known neuroanatomical structures. At three clusters, the features from both methods (AE and PCA) identified major tissue divisions (white matter, grey matter, borders & ventricles), while at nine and 21 clusters progressively detailed structures emerged (white matter variations, hippocampal subfields, and cortical layers).

AE-derived features demonstrated superior performance: more stable assignments across different cluster numbers and clearer anatomical boundaries, especially in cortex, hippocampus, and amygdala. This suggests that nonlinear feature learning better captures the intrinsic organizational properties of brain tissue, justifying the additional computational investment.

When AE-features from sham and mTBI animals were clustered together, the workflow identified pathology-associated signatures without prior injury information. A cluster that represented a small fraction of tissue in sham animals (5–6%) increased to 38.8% in one injured animal, capturing subtle tissue damage that may include axonal injury, inflammation, or glial changes. This suggests the ability to detect heterogeneous injury patterns across brain regions.

Recent work has shown that myelin organization patterns carry biological significance: myeloarchitectonic variations relate to functional network organization^33^, axonal morphological changes at the micrometer scale occur in neurological disorders^34,35^, and brain lipid distributions align with functional anatomy^36^. Our workflow demonstrates that unsupervised deep learning can automatically identify biologically meaningful tissue patterns and detect pathology without manual labeling, enabling scalable annotation-free approaches^13,20^.

This study has several limitations. Models were trained on a single staining batch from nine rat brains and have not been tested for generalization to different staining protocols or batches. While histogram matching addressed intensity variations, advanced staining normalization techniques developed for other stainings, such as H&E, could improve robustness if adapted for myelin staining^12^. The pathology-associated cluster identified in mTBI animals is intriguing but requires further characterization — histological and molecular markers could identify the cellular and biochemical composition to determine whether it represents inflammation, gliosis, axonal damage, or combinations thereof. Finally, our analysis treated tissue as 2D images, although myelin organization varies across the third dimension; automated 3D histology pipelines combining block-face imaging with whole-brain analysis^37^ could extend our workflow to volumetric tissue characterization.

In this work, we tested whether unsupervised deep learning could be used to quantify meaningful patterns in myelin-stained rat brain tissue sections without any labels. Our results show that convolutional AEs capture fine axonal organization details, and clustering these features recovers known neuroanatomical structures more effectively than linear methods, while also detecting pathology-associated features. These findings confirm our hypothesis that nonlinear unsupervised methods can overcome the limitations of traditional, object-centric histology workflows. By establishing a framework for label-free quantification of tissue architecture, our work addresses a gap in computational neuropathology and provides a foundation for future studies extending this approach to other staining methods, tissue types, or disease models.

## Resource availability

### Lead contact

Requests should be directed to the lead contact, Alejandra Sierra.

### Materials availability

This study did not generate new materials.

## Data and code availability

Our source code is available on GitHub (https://github.com/UEF-Multiscale-Imaging/AE-and-GMM-myelin-stained-tissue). Data used in this paper will be shared by the lead contact, Alejandra Sierra upon request. Requests are typically answered within 2 weeks.

## Acknowledgments

The authors wish to acknowledge CSC – IT Center for Science, Finland, and the UEF Bioinformatics Center (Biocenter Kuopio, Biocenter Finland, University of Eastern Finland) for computational resources, as well as the Biobank of Eastern Finland for section scans.

This work has been funded by the European Union’s Horizon 2020 research and innovation programme under the Marie Sk∤odowska-Curie agreement No 101034307, and by The Research Council of Finland via Flagship of Advanced Mathematics for Sensing Imaging and Modelling (FAME; grant number 358944).

## Author contributions

Conceptualization: ME, RS, VK, JT, AS; methodology: ME, RS, VK, JT, AS; software, validation, formal analysis and investigation: ME; resources: RS, ISMM, VK, JT, AS; data curation and writing - original draft: ME; writing - review & editing: all; visualization: ME, ON; supervision: RS, VK, JT, AS; project administration and funding acquisition: VK, JT, AS.

## Declaration of generative AI and AI-assisted technologies in the writing process

During the preparation of this work, the authors used Claude to help with refining language, clarity and organizing sections. After using this tool, the authors reviewed and edited the content as needed and take full responsibility for the content of the published article.

## Materials and methods

### Animals and histology

We used photomicrographs of 13 adult male Sprague-Dawley rats (10 weeks old, 300–450 g, Harlan Netherlands B.V) from a previous study^38^. Seven animals underwent mild traumatic brain injury (mTBI) induced using the lateral fluid percussion (LFP) injury model^39^. Briefly, under anesthesia, a 5mm craniectomy was performed on the convexity of the left skull. An LFP injury induced a mild brain injury (0.93 *±* 0.22 atm). Six other animals (shams) underwent the same surgical procedure except for the injury, serving as sham-operated controls. All experimental procedures were approved by the Animal Ethics Committee of the Southern Finland Provincial Government and conducted in accordance with the guidelines set by the European Union Directives 2010/63/EU.

Thirty-five days after surgery, all rats were anesthetized and transcardially perfused 4% paraformaldehyde (more details in ^38^). After perfusion, the brains were removed from the skull, postfixed, and then placed in a cryoprotective solution and frozen to store at −70^*°*^*C* until sectioning. All brains were sectioned in the coronal plane (30 *µm*, 1-in-5 series), using a sliding microtome. The data used in this study corresponds to one series stained with gold chloride for myelin^38,40^ to assess myeloarchitecture. After staining, high-resolution photomicrographs of the myelin-stained sections were acquired using a Hamamatsu scanner, NanoZoomerXR. The images were acquired at 40x, with a resolution of 226*nm/px*. Each photomicrograph corresponds to one microscope slide that contains typically six brain sections (Figure 6a) with the same staining.

**Figure 6:**
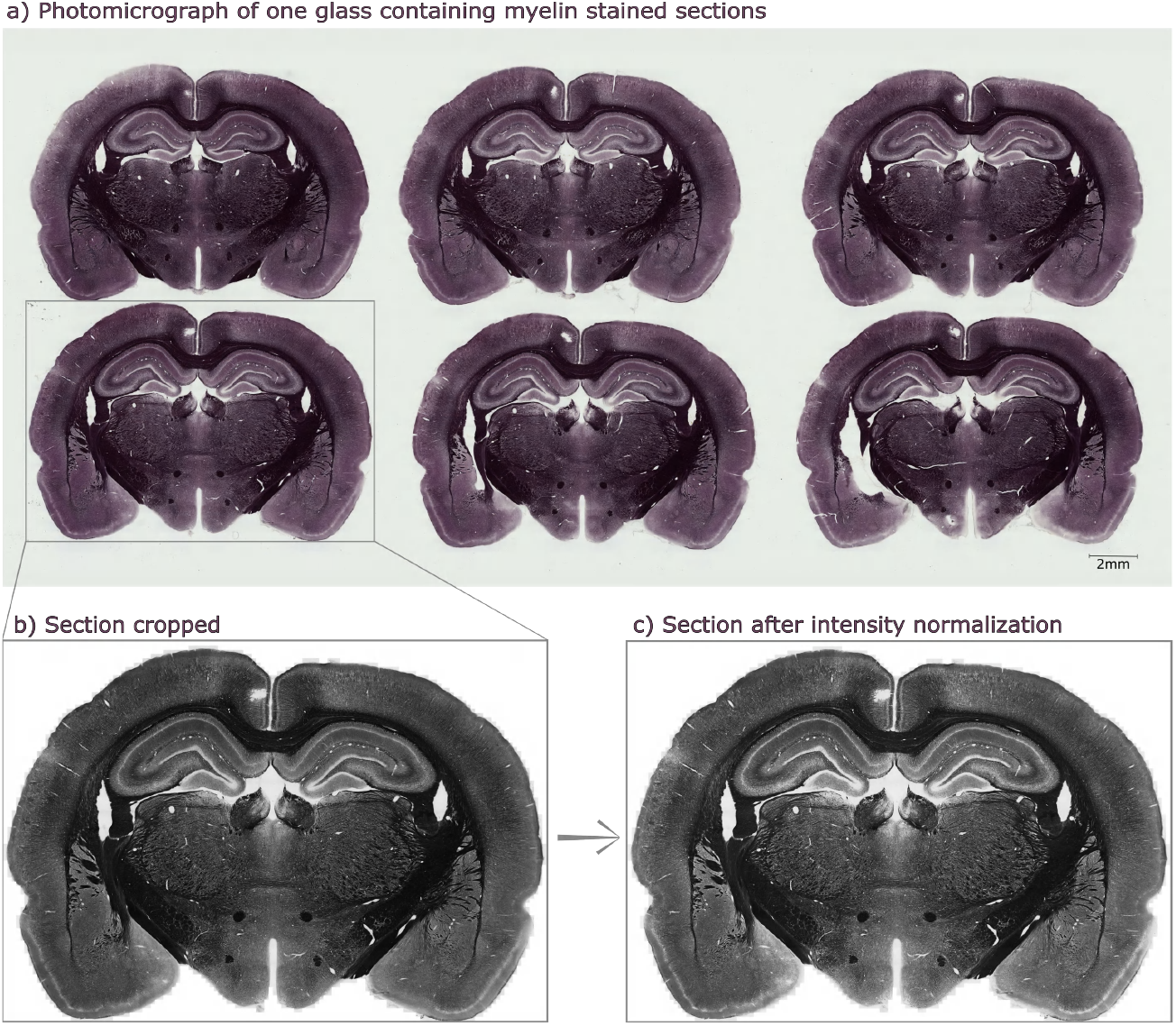
Preprocessing: a) photomicrograph of a glass containing six myelin-stained sections; b) one section after cropping; c) same section after intensity normalization.

### Image preprocessing

Image preprocessing consisted of first separating the photomicrographs of glasses into individual image files containing only one section; and second correcting for intensity staining variations across the sections. We preprocessed photomicrographs of sections +2.28 mm to -7.56 mm from bregma^29^. This selection resulted in a total of 592 photomicrographs.

We converted the photomicrographs to grayscale, downsampled them by a factor of 2^6^, and created a mask for each section. The masks served to define the coordinates of a bounding box containing each section. These coordinates were then rescaled to the original image size and used to crop each section (Figure 6b), which was saved in grayscale at the original resolution. As a result, we obtained individual images for each section.

We performed histogram matching^41^ to correct for staining-intensity variations across sections. For each rat brain, we used all image sections and their corresponding masks to remove background pixels, and selected one reference section from the central five slices for intensity normalization. An example of intensity normalization is shown in Figure 6c. We performed intensity normalization in three steps. 1) Within each brain, we performed iterative histogram matching in two directions (rostral and caudal), starting from the middle reference section. The reference section was histogram matched with its immediate neighbors - one section rostrally and one section caudally. The corrected sections then served as references for their respective next neighbors, with the process continuing iteratively in both rostral and caudal directions until all sections of the brain were processed. 2) We normalized across different brains by selecting one brain from a sham-operated rat as the global reference and histogram matching reference sections of all other brains to it. 3) We repeated the first step using the updated reference sections from step two, propagating the cross-brain normalization throughout all sections of the brain.

### Feature extraction

Of the 13 animals included in this study, 2 sham-operated and 2 mTBI animals animals were reserved for independent evaluation as a test set. The 2 mTBI animals in this group represented different pathological profiles -one with clear structural damage and the other with subtle injury. From the remaining 9 animals, one mTBI brain was randomly selected as the validation set (for the AE, not used in PCA). The remaining 8 brains (4 sham-operated and 4 mTBI) were used for training AE and PCA models. To minimize animal-specific bias, sections from all animals in the training set were shuffled before training AE and PCA. For both PCA and AE, we trained two separate models with different input patch sizes (128×128 pixels and 256×256 pixels), extracting 256 features from each patch. The 128×128 pixel size was chosen as the smallest patch that preserved sufficient histological information, while the patches of 256×256 pixels enabled comparison of model performance across spatial scales. The background patches were excluded from training. The number of training patches per section depended on the input patch size. For input patch size of 128×128 pixels, we extracted 102,400 non-overlapping patches per section, and for input patch size of 256×256 pixels we worked with 25,600 non-overlapping patches per section.

### Principal component analysis (PCA)

Given the large size of the dataset, we applied Incremental PCA^42^, which processes data in batches to enable memory-efficient decomposition. It was implemented using scikit-learn IncrementalPCA, with a batch size of 1024 samples and 256 components.

### Autoencoders (AE)

We used a convolutional AE architecture^27^ (Figure 7) to extract features from tissue patches. Regardless of the input patch size, the encoder’s first convolutional layer extracts 8 channels from the input patch while decreasing by half the patch size. The consecutive convolutional layers duplicate the number of channels, while decreasing by half the patch size. We used as many layers as necessary to reach a patch size of 1×1. Then, we flattened the vector and applied a linear layer to get the desired feature dimension. We applied batch normalization after each convolutional layer, followed by the ReLU activation function. The decoder was a mirror of the encoder architecture, except for a final sigmoid layer to ensure bounded output values in the range [0, 1]. All convolutional encoder and decoder layers used a kernel size of 3 and stride of 2.

**Figure 7:**
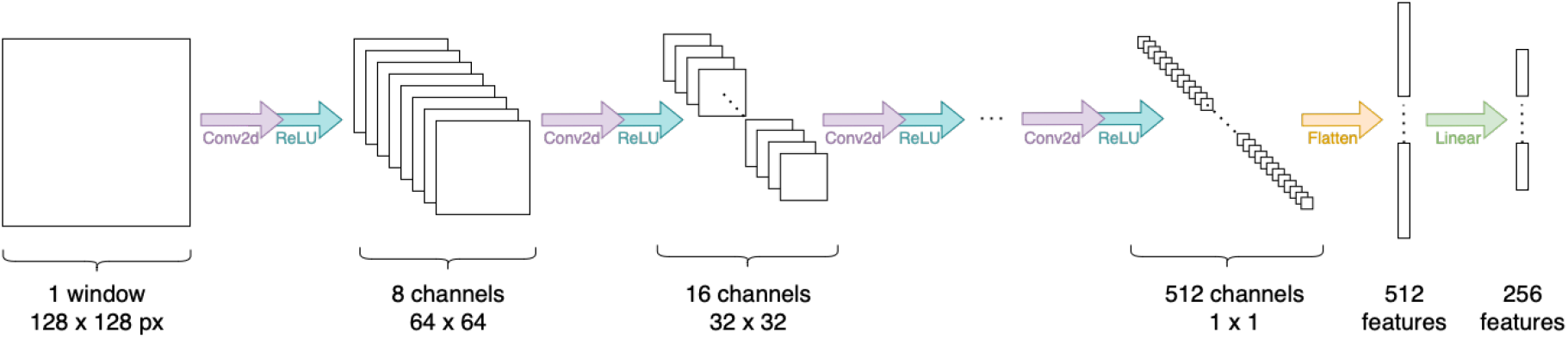
Diagram of the encoder architecture with input window size of 128×128 px and 256 features extracted.

**Figure 8:**
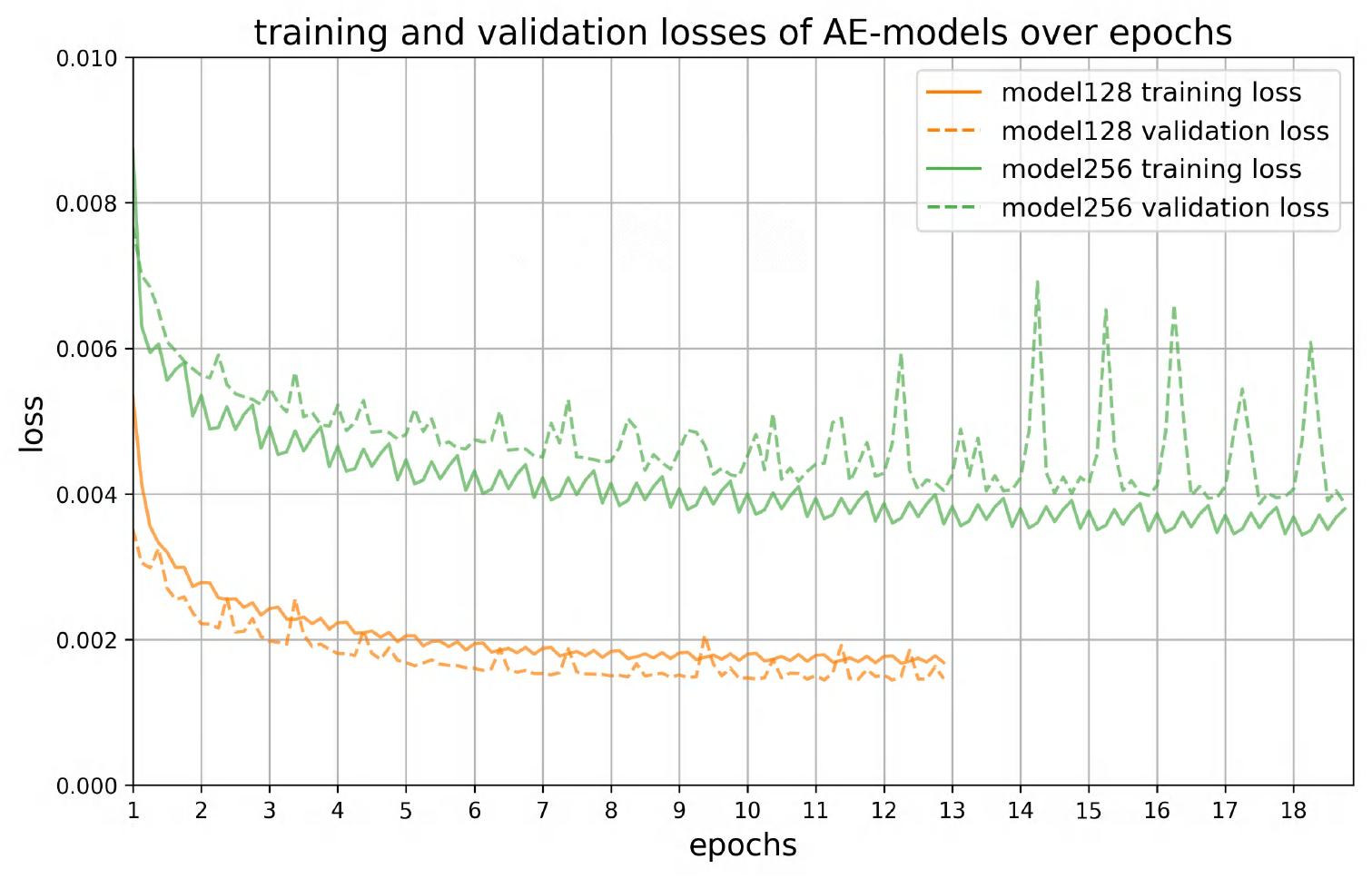
Training and validation loss curves for AE model128 and model256. Model128 shows stable convergence with training and validation losses remaining closely aligned. Model256 exhibits intermittent spikes in validation loss during the final third of training, suggesting potential instability when learning features from larger patches.

**Figure 9:**
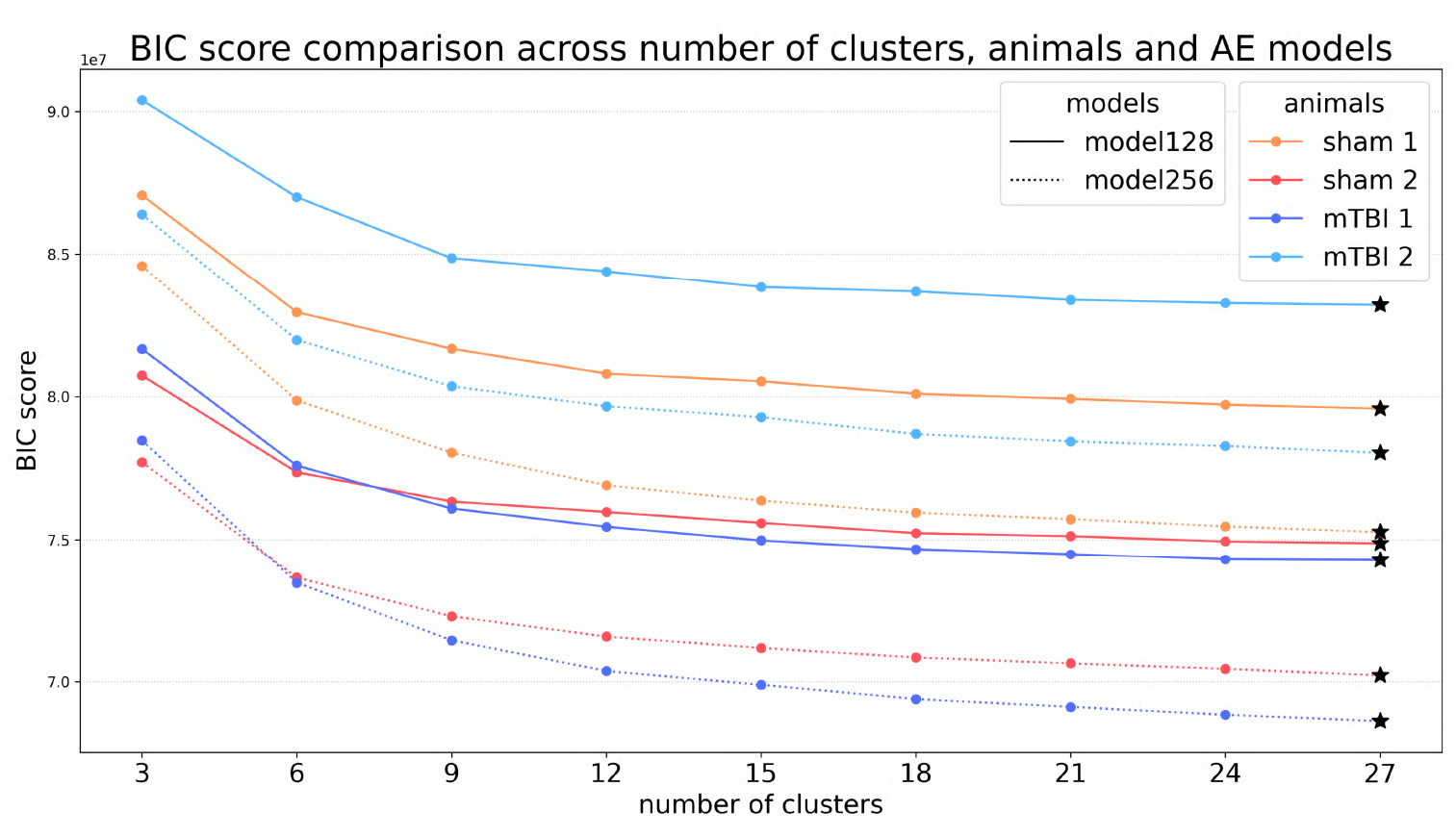
BIC scores from clustering AE-features. Regardless of the model or animal, the greater the number of clusters, the lower the BIC score, achieving the lowest (star symbol) at 27 clusters in this cluster range. The elbow is considered at 9 clusters, when the BIC score starts to stabilize.

**Figure 10:**
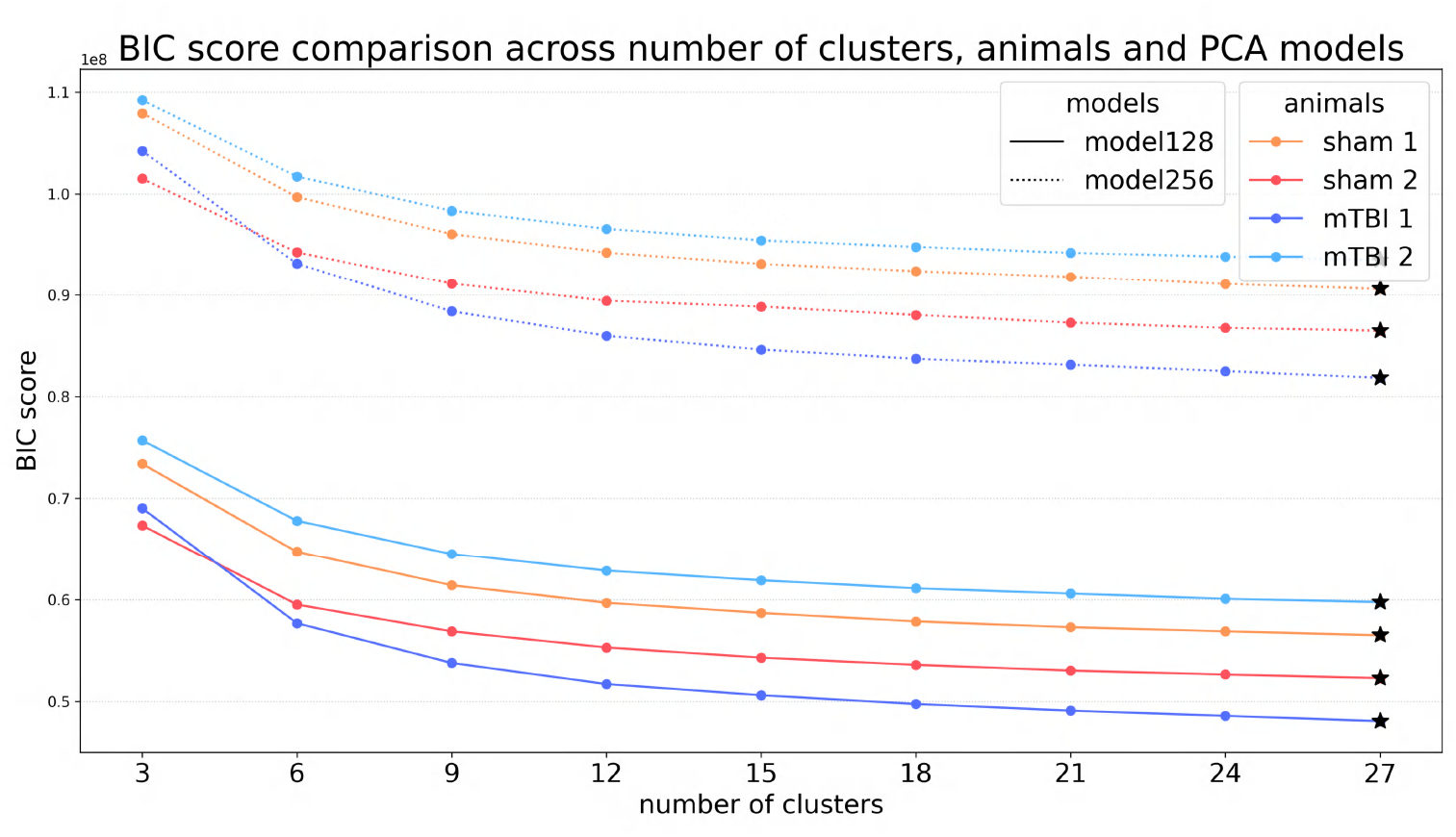
BIC scores from clustering PCA-features. Regardless of the model or animal, the greater the number of clusters, the lower the BIC score, achieving the lowest (star symbol) at 27 clusters in this cluster range. The elbow is considered at 9 clusters, when the BIC score starts to stabilize.

**Figure 11:**
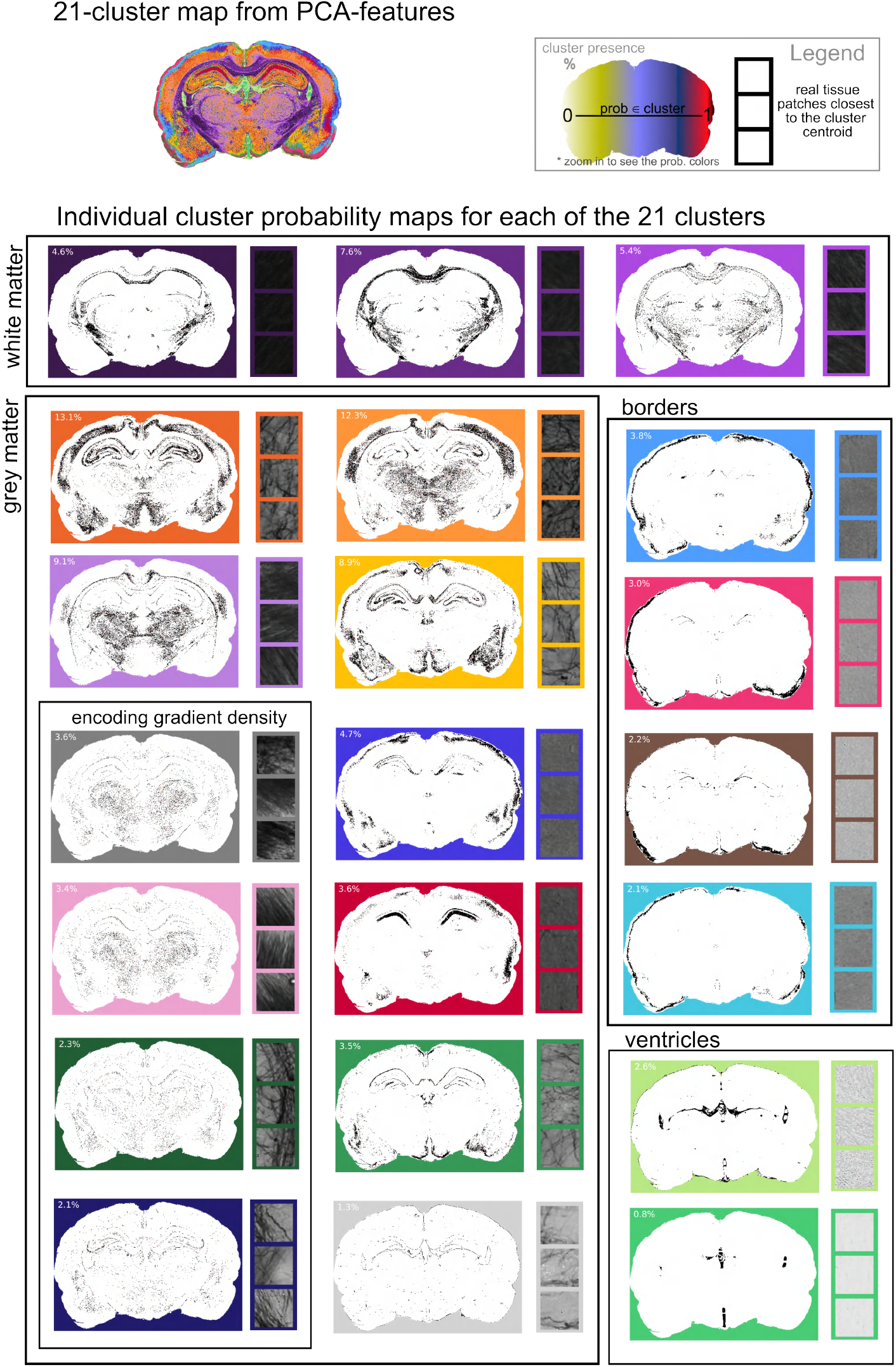
Probability maps of a single section. On the top left, overview of the 21-cluster map from PCA-features from one section. On top right, the legend describing the color scale of the probability maps. Then, the individual cluster probability maps grouped in: white matter, grey matter, borders and ventricles. Each probability map shows a percentage that indicates its presence in the tissue section, as well as three real tissue patches (128×128 pixels) closest to that cluster centroid.

We trained the model using MSE loss function and Adam optimizer with a learning rate of 10^−3^. The model was trained in batches of 64 windows. To prevent overfitting, we implemented early stopping with a patience of 10, meaning that training was halted if validation loss failed to improve for 10 consecutive evaluations. Validation loss was assessed periodically throughout each epoch, after processing every 1/8 of the training data. The best-performing state of the model was saved before patience counting began.

The dataset was processed dynamically, with patches extracted on demand rather than stored in memory. We combined Pytorch DataLoader with a window extraction function to handle this process. The training pipeline was implemented in Pytorch and executed on GPU resources from Puhti supercomputer at CSC – IT Center for Science, Finland. A single NVIDIA V100 GPU was allocated for training, along with 80 GB of memory.

### Feature analysis

#### Reconstruction quality assessment

To quantify reconstruction quality, we employed two complementary metrics: MSE and SSIM^30^.

MSE quantifies pixel-wise intensity differences between original and reconstructed images. For images *I* (original) and *R* (reconstructed), the MSE of the pair (*I, R*) was computed only on tissue pixels (i.e. where the mask > 0) as follows:

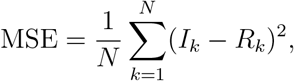

where *N* is the total number of tissue pixels within the mask.

SSIM measures perceptual similarity by comparing local patterns of luminance, contrast, and structure, providing complementary information to MSE about the preservation of spatial features. For images *I* and *R* normalized to [0,1], SSIM is defined as:

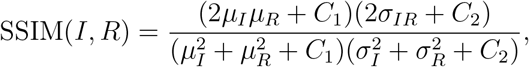

where *µ*_*I*_, *µ*_*R*_ are mean intensities; *σ*_*I*_, *σ*_*R*_ are the standard deviations; *σ*_*IR*_ is the covariance; and *C*_1_ = (*k*_1_*L*)^2^, *C*_2_ = (*k*_2_*L*)^2^ are stabilization constants with *k*_1_ = 0.01, *k*_2_ = 0.03, and *L* the dynamic range of pixel values^30^. Given the large dimensions of the whole-brain images, we computed SSIM on non-overlapping patches of 2048×2048 pixels, which were then averaged across all tissue-containing patches per section (defined as patches with more than 10% of tissue based on brain mask). This approach maintains the sensitivity of SSIM on local structural features while enabling efficient computation.

The values shown in Table 1 and Table 2 represent the mean of the MSE and SSIM values per animal obtained from its sections.

To assess the practical significance of reconstruction quality differences between PCA and AE, we computed Cohen’s d^43^, which standardizes the difference between group means by their pooled standard deviation:

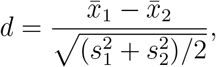

where 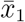,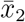 are group means and *s*_1_, *s*_2_ are standard deviations. This metric quantifies how many standard deviations separate the two methods. Effect sizes are conventionally interpreted as small (|*d*| ≈ 0.2), medium (|*d*| ≈ 0.5), or large (|*d*| ≈ 0.8)^43^.

#### Unsupervised clustering

GMM is a probabilistic clustering approach that models data as a weighted sum of multiple Gaussian distributions^28^. Each cluster is characterized by a mean and covariance matrix. We fit GMM using the Expectation-Maximization (EM) algorithm, which iteratively estimates cluster parameters and assigns probabilistic memberships to data points^44^.

After parameter exploration, we configured GMM with spherical covariance (independent variance per component) and covariance regularization of 10^−3^. Model parameters were initialized using k-means clustering and a regularization parameter of 10^−3^. For each animal, we randomly selected 100,000 tissue patches, extracted 256-dimensional features via PCA or AE encoders, and fit GMM on these feature representations.

We evaluated cluster numbers from 3 to 27 (in steps of 3, Supplementary Figure 9 and Supplementary Figure 10) using BIC^31^ to balance model fit with complexity. We selected three numbers of clusters for detailed analysis: 3 clusters to represent major tissue types; 9 clusters, corresponding to the elbow in the BIC curve where improvement plateaued across all animals; and 21 clusters to reveal finer tissue details while maintaining interpretability.

#### Cluster matching via symmetric KL divergence

To enable visual comparison across different numbers of clusters, we manually assigned colors to the 21-cluster solution, then sequentially propagated these colors to solutions with fewer clusters (18 down to 3, in steps of 3). For each propagation step, we computed the symmetric Kullback-Leibler (KL) divergence between all pairs of cluster distributions, modeled as spherical Gaussian distributions (multivariate normal with spherical covariance). We used the Hungarian algorithm to find the optimal one-to-one cluster assignment that minimized the total symmetric KL divergence, with each cluster inheriting the color of its optimal match.

Assuming two distributions *P ∼ 𝒩* (*µ*_0_, Σ_0_), *Q ∼ 𝒩* (*µ*_1_, Σ_1_) of dimension *k*, the symmetric KL divergence from *Q* to *P* is defined as

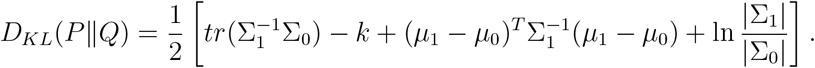

Let *I* be identity matrix of rank *k*. For spherical multivariate normal distributions, 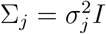 and therefore 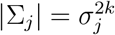 and 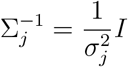 for *j* = 0, 1. Therefore, symmetrized KL divergence is:

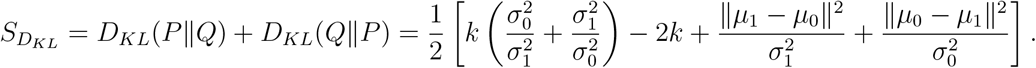

#### Probability maps and real tissue cluster representatives

For each tissue patch, the GMM model outputs probabilities for all clusters. From each tissue section, we generated spatial probability maps for every cluster, visualizing both the distribution and the confidence of the assignments. We also identified feature vectors nearest to each cluster centroid and mapped them back to the original patches, providing interpretable examples of what each cluster represents in the image space.

#### Cross-animal clustering

To assess the consistency and generalizability of cluster organization across animals, we performed clustering on a combined dataset. From each of the 4 animals used for evaluation, we randomly selected 25,000 tissue patches (excluding background), yielding a total of 100,000 patches from the 4 different animals. We applied the GMM on this dataset, and compared the cluster distribution across animals.

